# From low to high transmission: Diversity-dependent responses of *Plasmodium falciparum* population structure to transmission intensity

**DOI:** 10.64898/2026.04.07.717068

**Authors:** David Suarez-Salazar, Vladimir Corredor-Espinel, Mauricio Santos-Vega

**Affiliations:** Interdisciplinary Graduate Program in Genetics and Genomics, Texas A&M University, College Station, Texas, USA; Department of Entomology, Texas A&M University, College Station, Texas, USA; Mathematical and Computational Biology Research Group (BIOMAC), Universidad de los Andes, Bogotá, Colombia; Department of Public Health, Faculty of Medicine, Universidad Nacional de Colombia, Bogotá, Colombia; Department of Biological Sciences, Universidad de los Andes, Bogotá, Colombia

## Abstract

Genetic surveillance is increasingly used to track malaria transmission, yet genomic metrics can respond nonlinearly to changes in transmission intensity and depend on the diversity already present in the parasite population. Here, we present a stochastic agent-based model of hu-man–mosquito transmission that integrates SEIS-like epidemiological dynamics with within-host *Plasmodium falciparum* haplotype dynamics. By varying the maximum mosquito biting rate and the initial parasite diversity, we examine how transmission intensity and standing diversity jointly shape mixed infections, recombination, and long-term population structure across a continuous transmission gradient. Our study revealed a sequential pattern in which increasing biting intensity first increases infection prevalence and multiplicity of infection, then expands opportunities for outcrossing, and only thereafter increases effective recombination and recombinant haplotype generation. These responses are strongest in low– to intermediate transmission and tend to plateau at higher transmission levels. Initial population diversity constrains the amount of diversity that can be maintained and the magnitude of recombination output, while temporal trajectories show that haplotype evenness can pass through transient non-equilibrium phases before stabilizing. Together, these results show that the structure of the parasite population is shaped not by transmission intensity alone but by its interaction with standing genetic diversity. Furthermore, this study works to clarify when and how genomic metrics reliably reflect transmission conditions across heterogeneous malaria settings.

## Introduction

Despite increased control efforts, malaria remains a major global health challenge, with the heaviest burden centered in sub-Saharan Africa (SSA) [1]. The World Malaria Report 2024 estimates 263 million malaria cases worldwide in 2023 and that the WHO African Region accounted for 94% of global cases and 95% of malaria deaths that year [1]. Among human malaria parasites, *Plasmodium falciparum* is responsible for the most severe disease and mortality, and its capacity to evade host immunity sustains transmission and complicates disease control efforts [2, 3, 4, 5].

Beyond mere case counts, *Plasmodium falciparum* populations exhibit significant genetic diversity and structure that influence transmission and shape the evolutionary trajectories of parasites in a given environment [6]. Genetic variation underpins immune evasion and facilitates the emergence and spread of drug-resistant lineages, meaning that similar epidemiological burdens can mask markedly different population-genetic states with distinct implications for intervention effectiveness [7, 8, 9]. As sequencing efforts expand, genetic surveillance has become an increasingly relevant method for interpreting transmission and connectivity [10, 11, 12, 13]. Unfortunately, mechanistic links between epidemiological conditions and emergent parasite structure remain difficult to isolate in field data where multiple drivers covary despite increases in sequencing efforts [14, 15].

Underlying difficulty in genetic surveillance methods lies in changes over populations due to the environment [15, 16]. Parasite population structure is not static: it emerges from transmission processes that determine the complexity of within-host infection, ranging from monoclonal to polyclonal infections, and thus the opportunities for genetic mixing and recombination [17, 18]. Sexual reproduction in the mosquito midgut involves meiosis and recombination, producing novel haplotypes whose establishment may depend on transmission frequency and the co-occurrence of different parasite genotypes within hosts [7, 19, 20]. Because recombination in *Plasmodium falciparum* is tightly linked to multiple-genotype infections, shifts in infection complexity can translate into nonlinear changes in diversity generation and relatedness at the population level [7, 21, 22, 23].

Transmission intensity is usually described using broad categories like high, moderate, or low based on epidemiological metrics that vary significantly both between and within endemic countries, intermediate regimes are less well understood [24, 25]. Capturing this heterogeneity requires a framework that treats intensity as a continuum and allows comparable evaluation across regimes [26, 27, 11]. A natural way to operationalize transmission intensity is through the force of infection, which can be modulated by the mosquito biting rate and is constrained by host–vector contact structure [28, 29, 30, 31].

Initial parasite genetic diversity constitutes an important boundary condition for these dynamics because it defines the amount of variation available for recombination and reshaping by transmission over time [32, 8, 33]. In real systems, transmission intensity and diversity are often coupled through demography, migration, interventions, and sampling, making it challenging to attribute observed shifts in genetic structure to a specific mechanism [14, 15, 32]. Disentangling the effects of transmission intensity from those of initial diversity, therefore, is essential for developing mechanistic expectations that can inform the interpretation of genetic surveillance patterns across the range of transmission settings [11, 26].

Here, we present a stochastic model that links malaria transmission to changes in parasite genetics, enabling controlled investigation of how evolutionary paths develop across varying transmission levels. By adjusting the biting rate, which influences infection force, and initial parasite diversity, we examine how both epidemiological factors and genetic starting points affect infection complexity within hosts and the overall population structure. Outcomes are assessed using various epidemiological and genetic metrics, such as the prevalence of monoclonal and polyclonal infections, multiplicity of infection, haplotypic combinations that are new to the circulating population, thereby measuring the conversion of coinfection into population-level genetic innovation. Special focus is given to understanding nonlinear behaviors and regime changes at moderate transmission levels.

## Methods

### Model Description

A stochastic, agent-based model that combines human and mosquito populations was developed to simulate malaria transmission and parasite genetic changes under controlled conditions of transmission intensity and initial genetic diversity. The overall model structure and simulation setup are summarized in Figure 1. An expanded view of the human and mosquito state-transition structure, including monoclonal, polyclonal, and mixed-stage infection classes, is provided in Supplementary Section S1 and Figure S1.

**Figure 1.**
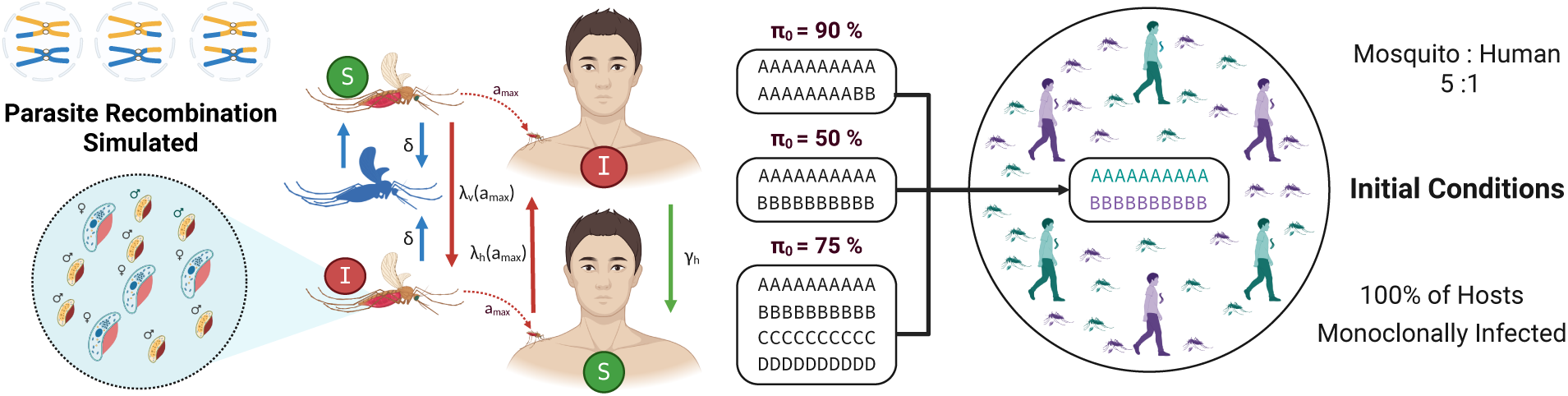
Model structure and simulation setup. Schematic representation of the transmission model and initial conditions used in the simulations. Parasite recombination is simulated within the mosquito vector, where sexual stages allow the generation of recombinant haplotypes. The epidemiological dynamics follow a vector–host transmission cycle between mosquitoes and humans, with susceptible (S) and infected (I) host states and bidirectional transmission between them. Different levels of initial parasite genetic diversity (*π*_0_) are explored, illustrated through example haplotype compositions. Simulations start with monoclonally infected hosts and vectors, each carrying a single haplotype. The initial population is configured with a mosquito-to-human ratio of 5:1. Colors within the population illustrate distinct parasite haplotypes circulating in the system. Created in BioRender. Suarez Salazar, C. (2026) https://BioRender.com/f920yd8

The epidemiological component utilized the SEIS vector–host transmission model outlined by Chitnis et al. 2008 [28]. This model explains how humans become susceptible to infection after being bitten by infectious mosquitoes, while mosquitoes acquire infection by feeding on infectious humans carrying transmissible parasite stages. After an extrinsic development period within the mosquito, infected mosquitoes can transmit the parasite to humans. Infections in both populations were classified as monoclonal or polyclonal based on the number of haplotypes carried by each individual. Multiplicity of infection served as a population-level indicator of infection complexity.

Meanwhile, the genetic component models parasite populations as haplotypes, recombining during the mosquito stage and thus connecting transmission dynamics with shifts in parasite population structure. In the simulations presented here, each haplotype consisted of 75 positions, which was sufficient to ensure that the number of observed haplotypes did not reach saturation. The initial parasite diversity was drawn from a predefined haplotype pool, distributed among infected hosts and vectors at the start of each simulation, enabling the model to analyze how the original genetic makeup influences subsequent transmission and recombination. Further details on parasite representation and haplotype diversity initialization are available in Supplementary Section S4.

### Model Formulation

The transmission process was formalized in terms of host– and vector-specific forces of infection, which determine the rates at which susceptible humans and mosquitoes acquire infection.

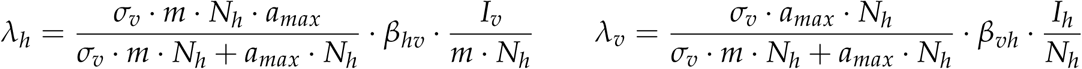

Following the framework of Chitnis et al. 2008 [28], the human availability term, previously denoted as *σ_h_*, is here expressed as *a_max_*for clarity, representing the maximum number of mosquito bites a human can receive per unit time. In this formulation, *N_h_* denotes the total number of humans, *N_v_* = *mN_h_* the total number of mosquitoes, *m* the mosquito-to-human ratio, *σ_v_*the mosquito biting demand, *β_hv_* and *β_vh_* the transmission probabilities from mosquito to human and from human to mosquito, respectively. A full derivation of the host and vector-specific force-of-infection terms is provided in Supplementary Section S2, and baseline parameter definitions, explored ranges, and literature sources are summarized in Table S1. Human and mosquito states were updated through a combination of stochastic infection-related events and individual-level developmental timers. Additional details on event scheduling, developmental timers, and overlapping infections are provided in Supplementary Section S3.

Specifically, infection of humans and mosquitoes, human recovery (*γ_h_*), and mosquito death (*δ*) were treated stochastically, whereas parasite development within hosts and vectors, including gametocyte maturation in humans (*α_h_*) and sporozoite maturation in mosquitoes (*α_v_*), was represented through deterministic timers assigned at the individual level. Parasite populations were represented as haplotypes that could accumulate in hosts and vectors, enabling repeated, polyclonal infections. When a mosquito acquires multiple haplotypes during blood feeding, recombination could occur during the sexual stage, generating recombinant offspring and linking transmission dynamics to changes in parasite population structure. The mosquito-stage recombination procedure is described in Supplementary Section S5. Some implementation components were adapted from the open-source SEIRS+ framework [34].

### Model Analyses and Simulations

To evaluate how transmission intensity and standing parasite diversity jointly shaped epidemiological and genetic outcomes, we simulated a factorial set of scenarios spanning five levels of initial genetic diversity (*π*_0_ = 10%, 30%, 50%, 75%, and 90%) and a continuous gradient of transmission intensity controlled through *a_max_* from 0.1 bites/day to 3 bites/day. At initialization, all humans and mosquitoes were initially infected, each carrying a single haplotype to ensure an even distribution of haplotypes across the population. Each scenario was simulated over eight years with 30 independent replicates, allowing the system to evolve from its initial conditions toward a steady state. Our analysis focused on the stable phase of the simulations, during which we measured infection dynamics and parasite population structure. Epidemiologically, we evaluated the overall prevalence of human infection, the proportions of monoclonal and polyclonal infections in humans and mosquitoes, the force of infection, and the multiplicity of infection in both hosts. Genetically, we examined changes in haplotype diversity, normalized Shannon index, haplotype count, and an effective recombination ratio — the fraction of recombination events that generated haplotypes absent from the circulating parasite population at that time. Detailed definitions of all epidemiological and genetic metrics are available in Supplementary Section S6. Additional simulations varying human population size and vector-to-host ratio were conducted to test the robustness of these patterns across different demographic scales (Supplementary Section S8). The code used to implement the model and reproduce the simulations is publicly available at https://github.com/dsuarez-s/PlasmoEpiGen.

## Results

Figure 2 illustrates how the multiplicity of infection (MOI), the effective recombination ratio (ERR), and the number of recombinant haplotypes (NRH) jointly vary across changes in maximum biting rate (*a_max_*) and initial parasite genetic diversity (I.G.D., *π*_0_). This broad range is consistent with the transmission spectrum shown in Figure S3, where the mosquito force of infection spans a wide range and increases considerably across the same parameter combinations.

**Figure 2.**
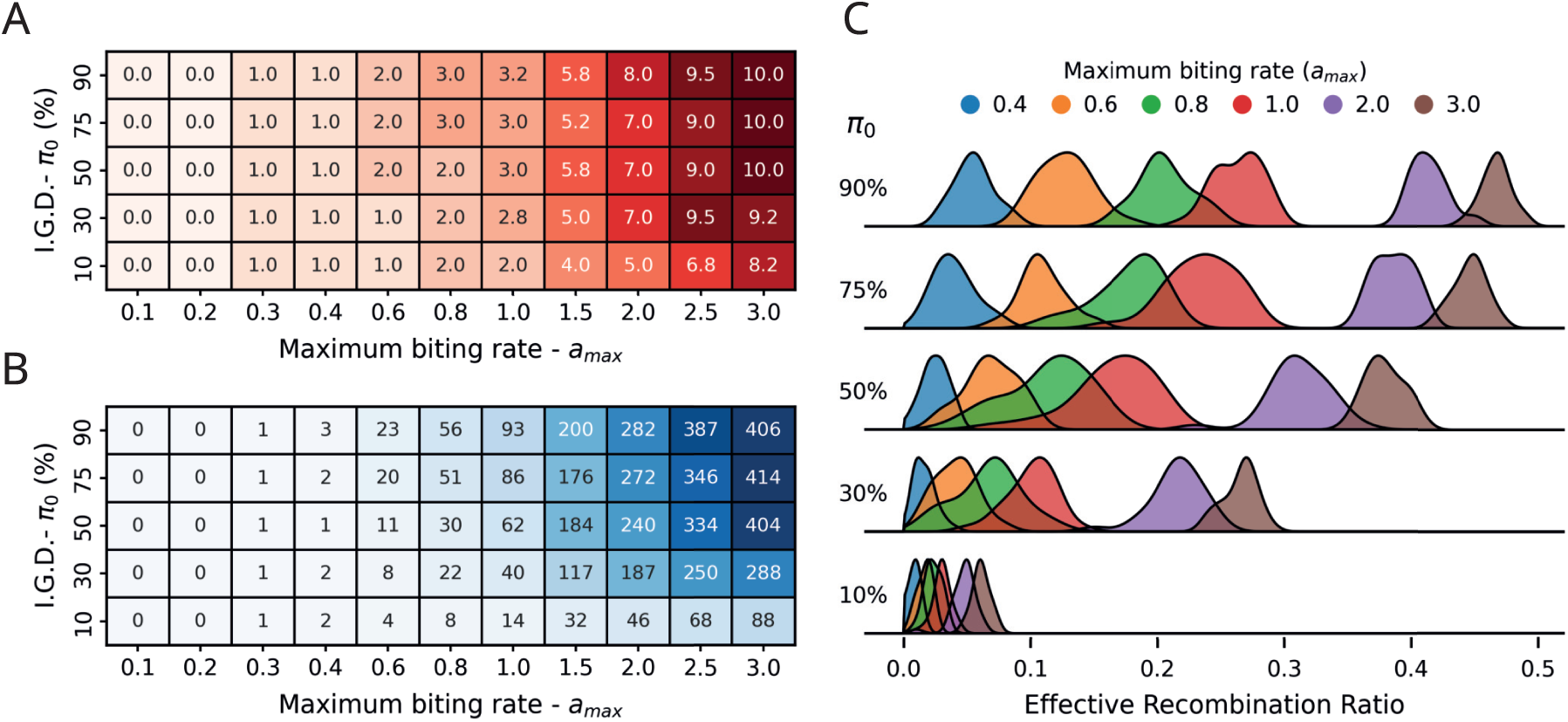
Human multiplicity of infection, circulating haplotypes, and the effective recombination ratio at the system steady state. (A) Heatmap of the median multiplicity of infection (MOI) in the human host population. (B) Heatmap of the median number of circulating recombinant haplotypes (NRH), representing the total diversity maintained in the system. (C) Ridgeline distributions of the effective recombination ratio, defined as a system-level measure of recombinant novelty generation. Specifically, it is computed as the fraction of all infection events that give rise to novel recombinant variants, and thus captures the overall contribution of recombination to haplotypic innovation rather than the outcome of individual recombination events. Complementary steady-state epidemiological responses across the same transmission gradient, including infection prevalence, polyclonal infection prevalence, force of infection, and multiplicity of infection in hu-mans and mosquitoes, are shown in Figures S2 and S3. Additional steady-state profiles for the effective recombination ratio and total haplotype richness are provided in Figure S5.

Panel A shows that the MOI remains at or near zero across all levels of initial diversity when *a_max_*is low (≤ 0.4), indicating predominantly monoclonal infections in human hosts at minimal transmission intensity, with little to no opportunity for recombination. As *a_max_*increases, the recombination scenario becomes possible, allowing two or more parasite genotypes to coexist within the same host. Beyond 1.0, MOI rises substantially, reaching up to 10 coinfecting haplotypes at the highest biting rates and diversity levels, demonstrating a clear and consistent positive relationship between transmission intensity and MOI. Panel B reveals a similar pattern in the absolute number of recombinant haplotypes generated, which increases from near zero at low biting rates to over 400 unique haplotypes at *a_max_*= 3.0 and *π*_0_ = 90%, highlighting the combined effect of high transmission and high initial diversity. Something important to note is the saturation effect observed in both MOI and NRH, where further increases in I.G.D. no longer produce proportional increases once transmission is already high. Together, Panels A and B suggest that MOI links transmission intensity to recombination output: higher biting rates increase coinfection within hosts, while higher initial diversity determines how much of that coinfection is translated into recombinant haplotypes.

Panel C further decomposes these dynamics by showing the full distribution of ERR across biting-rate levels for each initial-diversity level. At low *π*_0_ (10%), distributions are tightly clustered near zero regardless of *a_max_*, whereas at higher diversity levels (50–90%), distributions broaden markedly and shift rightward with increasing biting rate, reflecting greater heterogeneity in recombination outcomes. Collectively, these results demonstrate that transmission intensity and initial genetic diversity interact to determine the magnitude and variability of recombination output, with important implications for how population genetic structure is shaped across different transmission settings.

This pattern is also consistent with Figure S2, where infection prevalence begins to saturate at *a_max_*values above approximately 1.5, and the proportion of polyclonal infections shows a similar saturation trend. In both cases, saturation occurs when more than 80% of humans are infected and, among those infected, more than 70% of infections are polyclonal. Interestingly, even under these highly transmissible conditions, a fraction of infections remains monoclonal, suggesting that increased transmission does not completely eliminate heterogeneity in within-host infections. This may also help explain why MOI and NRH in Figure 2 begin to level off at high *a_max_*values, since once infection prevalence and polyclonality are already high, further increases in transmission may yield diminishing returns.

Population genetic structure is characterized here using two indices: haplotypic genetic diversity and the normalized Shannon index. Together, these metrics capture both the richness of haplotypes and their evenness across hosts. Figure 3 examines the down-stream consequences of transmission-driven recombination for parasite population structure, quantified using two complementary metrics: the loss of genetic diversity relative to initial conditions (Panel A) and the evenness of the resulting haplotype distribution, measured by the normalized Shannon Index (Panel B).

**Figure 3.**
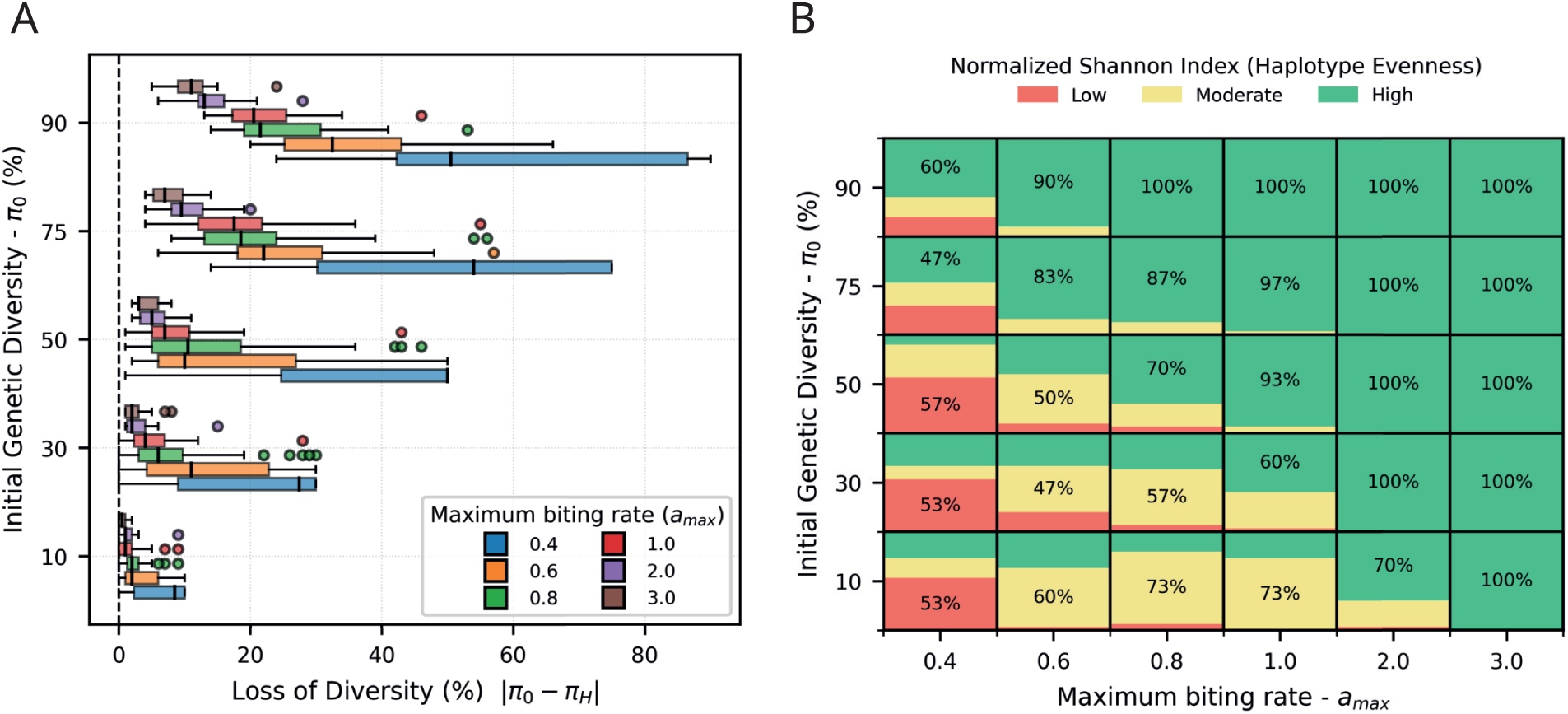
Genetic diversity loss and haplotype evenness at steady state. (A) Boxplots show the loss of human haplotypic genetic diversity as the difference between the initial diversity level and the steady state diversity level. Results are grouped by the initial genetic diversity level on the y axis and colored by the maximum biting rate. (B) A grid of stacked proportions summarizes the steady state normalized Shannon index of haplotype evenness. Values are binned into low evenness from 0 to 0.3, moderate evenness from 0.3 to 0.7, and high evenness from 0.7 to 1. Each cell represents one combination of initial genetic diversity and maximum biting rate and partitions all simulations across the three bins. The number inside each cell reports the percentage of simulations in the most frequent bin. Related steady-state trends in haplotypic genetic diversity and normalized Shannon index across the same transmission gradient are shown in Figure S4.

Panel A demonstrates that diversity loss is greatest at low biting rates (*a_max_* ≤ 0.6) across all initial diversity levels, with several distributions extending toward high loss values, particularly at intermediate to high initial diversity levels (*π*_0_ = 30–90%), where median losses can exceed 20–50% and whiskers extend beyond 80%. However, at higher *a_max_* values, the parasite population retains more of its initial diversity, indicating that stronger transmission is associated with greater preservation of haplotypic variation. Notably, the relationship between initial diversity and diversity loss is not strictly monotonic: intermediate *π*_0_ values can sometimes exhibit losses comparable to or even greater than those under the highest-diversity conditions, suggesting nonlinear dynamics in how transmission and recombination reshape haplotype frequency structure.

Panel B complements these findings by revealing that haplotype evenness transitions sharply from predominantly low (red) to high (green) as amax increases, with the proportion of simulations achieving high evenness reaching 100% at *a_max_* ≥ 2.0 across nearly all initial diversity levels. At lower biting rates (*a_max_* = 0.4–0.6), a substantial fraction of simulations yield low or moderate evenness, reflecting the persistence of unequal haplotype frequencies under limited recombination and suggesting the presence of a dominant haplotype.

Together, these results indicate that increasing transmission intensity is associated with lower diversity loss while simultaneously homogenising the haplotype distribution, producing populations that retain more genetic diversity overall and are more evenly structured. Conversely, low transmission is more consistent with the persistence of dominant haplotypes — a pattern that has important implications for interpreting genetic surveillance data across transmission settings.

The temporal dynamics of these steady-state regimes under low-, intermediate-, and high-transmission scenarios were further investigated, highlighting that recombination-and diversity-related metrics stabilize at different rates depending on biting intensity and initial genetic diversity (Figure S6). In particular, Figure S6 shows that under low transmission, recombination remains limited while haplotypic diversity and Shannon diversity decline progressively over time, consistent with the increasing dominance of one or a few haplotypes. Under intermediate-to-high transmission, the temporal dynamics of the Shannon index further suggest a transient bottleneck-like phase, in which evenness initially declines before recovering, consistent with the temporary dominance of one or a few haplotypes prior to the establishment of the final population structure. By contrast, as recombination rises more rapidly, the diversity metrics stabilize at lower rates.

Additional simulations varying human population size (*N_h_*= 30, 50, 70) and the mosquito-to-human ratio (*m* = 3, 5, 7) showed that the main transmission-genetic relationships remained qualitatively consistent across demographic settings (Supplementary Section S8; Figures S7-S14). Across these sensitivity runs, the mosquito-to-human ratio had a stronger effect than human population size on epidemiological outcomes, whereas haplotype richness and diversity metrics also retained a detectable dependence on system size, especially under intermediate-to-high transmission and moderate-to-high initial genetic diversity. These patterns are consistent with the contact formulation used here, because varying m changes the encounter structure itself, whereas varying *N_h_* at fixed *m* primarily changes the total number of transmission opportunities available to maintain and reshuffle haplotypes.

## Discussion

Our results support the idea that malaria transmission leaves detectable genomic signatures, but with one key refinement: transmission intensity alone does not determine these signatures. Parasite population structure results from the interaction of transmission intensity, infection complexity within hosts, and the diversity of haplotypes available for recombination. This explains why genomic metrics reliably track transmission in some contexts but can be unpredictable in others, especially when populations are not at equilibrium or when the historical makeup of circulating lineages varies across locations [26, 27, 35, 14].

A central contribution of this study is resolving the transition from low to high transmission as a continuum rather than a switch between discrete states. As biting intensity increases, infection prevalence and mixed infection rates rise first, expanding the opportunity for outcrossing, and only subsequently translating into a higher effective recombination ratio and greater haplotypic novelty. This sequential dynamic is consistent with modelling and empirical work showing that metrics linked to mixed infections respond strongly and rapidly to transmission intensity, on the order of months, while internal infection composition depends on the balance between cotransmission and superinfection [26, 27, 22, 23]. Critically, this progression plateaus at higher transmission levels: both epidemiological and recombination-related responses saturate as polyclonality becomes widespread. Transmission may continue to rise even as genetic indicators stall, a caution for surveillance programmes that rely on diversity metrics as proxies for burden [26, 27, 16].

Figure 3 demonstrates that long-term population effects go beyond just recombination rates. Low-transmission environments cause a gradual loss of diversity and decreased haplotype evenness, which aligns with the ongoing presence of dominant haplotypes in limited outcrossing scenarios. As transmission increases, populations preserve more diversity and become more balanced, showing recombination’s ability to oppose the merging of lineages into a few dominant backgrounds. Importantly, the initial standing diversity limits how far this process can go: transmission alone does not set the maximum level of haplotypic diversity [22, 8, 23, 32].

The temporal trajectories in Figure S6 further illustrate that these end states are reached through qualitatively different paths. Under low transmission, recombination remains limited while diversity and evenness decline gradually, consistent with the slow establishment of dominant haplotypes. Under intermediate and high transmission, however, evenness can initially decrease before recovering, indicating a transient bottleneck-like phase before the final population structure stabilizes. This point is important because it suggests that genetic observations taken during epidemiological change may not reflect equilibrium expectations, even if they eventually do. In that respect, our findings align with recent modeling studies showing that some malaria genetic metrics respond quickly to changes in prevalence, while others are slower, more variable, or highly dependent on the epidemiological driver of change [26, 27]. Overall, they emphasize the need to interpret genomic surveillance data in the context of transmission history and transition dynamics, not just final prevalence levels.

Our sensitivity analyses reveal that varying the vector-to-host ratio and varying host population size are mechanistically distinct perturbations (Supplementary Section S8). Varying *m* = *N_v_*/*N_h_* directly alters the per-capita encounter structure and thus the conditions that sustain mixed infection and recombination. By contrast, varying *N_h_*at fixed *m* primarily changes system size and the degree of stochastic haplotype retention. This distinction has direct implications for interpreting intervention-driven genomic change: vector control that reduces mosquito density relative to hosts is expected not only to suppress prevalence but to reduce polyclonality, effective recombination, and long-term haplotypic diversity — a compounded genomic consequence that demographic differences in host population size would not replicate.

Several limitations bound these conclusions. Transmission in our framework is driven through the maximum biting rate, making our findings strongest for vector-mediated change rather than changes arising from treatment, immunity, or migration. The model begins from controlled initial diversity conditions with monoclonal host infections — a deliberate simplification that isolates the mechanism but cannot capture the historical memory of natural parasite populations. We do not model importation, selection, spatial structure, or heterogeneous recombination landscapes. These omissions matter: low-transmission settings can retain substantial diversity when importation is frequent [36, 12, 37], and malaria genomic epidemiology is increasingly framed around effective recombination as the primary inferential quantity of interest rather than diversity alone [38]. Our results are best read as a mechanistic baseline for locally generated population structure rather than a complete account of all real-world settings.

## Conclusion

This study highlights broader questions about how transmission intensity and standing diversity together influence parasite population structure. Our findings clarify the interplay of mixed infections, recombination, and haplotype organization across different transmission levels. However, they also raise questions about how these relationships might change with additional biological processes. For example, it is still unclear whether the diversity seen in low– or intermediate-transmission areas is mainly generated locally through recombination or replenished by migration of new haplotypes [12, 37, 36]. It is also uncertain how non-uniform recombination landscapes could affect the assembly and stability of new genetic backgrounds [7, 39, 38], and how selection might spread adaptive or drug-resistant variants under various epidemiological conditions [9, 40, 41]. Addressing these issues is vital to determine when genomic patterns reflect local transmission alone versus the combined effects of migration, recombination, and selection. Ultimately, these results demonstrate that malaria parasite population structure depends not only on transmission intensity but also on its interaction with diversity and recombination, explaining why different genomic signatures can arise under similar epidemiological conditions.

## Supporting information

License Figure 1

License Figure S1

