## Supplementary material for "From low to high transmission: Diversity-dependent responses of *Plasmodium falciparum* population structure to transmission intensity": Suplementary Material

### Supplementary Information

#### S1. Extended Model Structure

Our model combines epidemiological and parasite genetic dynamics in a coupled human-mosquito system. Both humans and mosquitoes are represented at the individual level and can occupy susceptible, exposed, or infectious states, depending on whether they have acquired parasites and whether those parasites have matured to transmissible stages. To capture infection complexity, infectious states were further classified as monoclonal or polyclonal based on the number of haplotypes each individual carries.

In humans, susceptible individuals become exposed following inoculation by infectious mosquitoes. Exposed infections were classified as monoclonal or polyclonal depending on the composition of the inoculated parasite set. After parasite development within the human host, exposed infections transitioned to infectious states. Because new inoculation events may occur before an existing infection is cleared, infectious humans could also transiently carry newly acquired immature parasites from later exposure. Recovery from infection returns humans to the susceptible state.

Mosquitoes follow a similar process. Susceptible mosquitoes become exposed after feeding on an infectious human carrying transmissible parasites. Like humans, mosquitoes are classified as monoclonal or polyclonal based on the haplotypes they carry in exposed and infectious states. Once the parasite matures inside the mosquito, it can transmit it to new human hosts. This extended state structure allows the model to capture repeated exposure, overlapping infections, and evolving infection complexity over time, all while maintaining a clear distinction between epidemiological status and parasite genetic makeup.

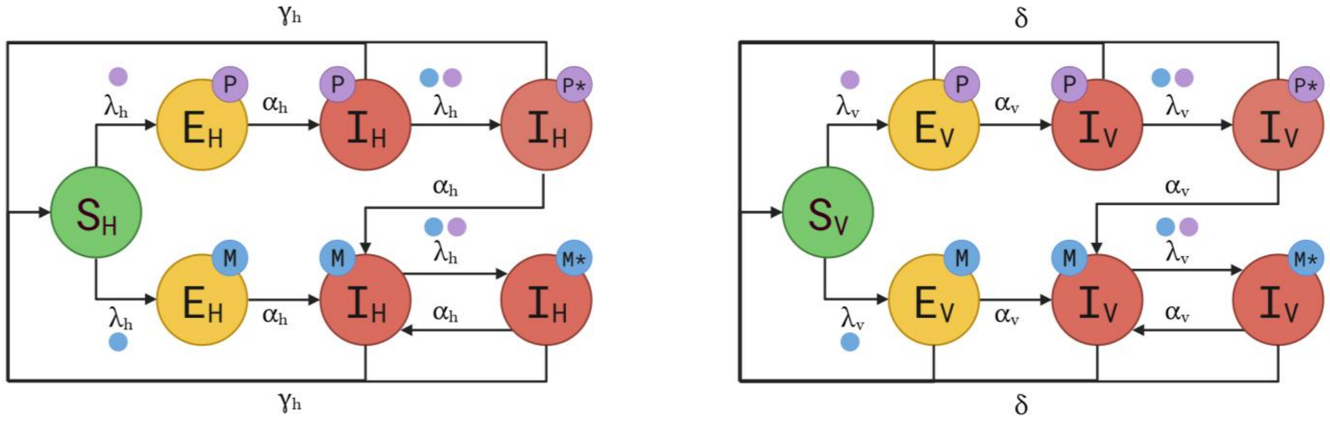


**Figure S1. Extended state-transition structure for human and mosquito infections.** The left panel shows the human submodel and the right panel the mosquito submodel. In each host type, susceptible individuals (S) can acquire either monoclonal or polyclonal infections, represented by the exposed (E) and infectious (I) states. Blue labels indicate monoclonal infections (M), whereas purple labels indicate polyclonal infections (P). States marked with an asterisk (M∗, P∗) denote individuals that are already infectious but have received an additional inoculation, so that mature parasites coexist with newly acquired immature parasites still undergoing development. After maturation, these mixed-stage states transition back into the corresponding infectious class without the asterisk. Created in BioRender. Suarez Salazar, C. (2026) https://BioRender.com/xxj2op8

#### S2. Force of infection formulation

Transmission between humans and mosquitoes was formalized in terms of human-specific and vector-specific forces of infection, $\lambda_h$ and $\lambda_v$. Following the mathematical framework (Chitnis, 2008), the human availability term, previously denoted as $\sigma_h$, is here written as $a_{max}$ to emphasize that it represents the maximum number of mosquito bites a human can receive per unit time. The mosquito biting demand is denoted by $\sigma_v$.

Let $N_h$ and $N_v$ be the total numbers of humans and mosquitoes, respectively. Then, $\sigma_v\, N_v$ represents the total number of bites required by mosquitoes per unit time to complete their gonotrophic cycles, while $a_{max}\, N_h$ indicates the total human biting availability in the population. Accordingly, the total number of effective encounters ($E_T$) between mosquitoes and humans per unit time is represented by half of the harmonic mean of the mosquito biting demand and human biting availability.

$$E_T = \frac{1}{\frac{1}{\sigma_v N_v} + \frac{1}{a_{max} N_h}} = \frac{a_{max}\, N_h\, \sigma_v\, N_v}{a_{max} N_h + \sigma_v N_v}\ bites \cdot day^{-1}$$

Here, $E_h$ represents the average number of effective encounters experienced by each human per unit time.

$$E_h = \frac{E_T}{N_h} = \frac{\sigma_v\, N_v\, a_{max}}{\sigma_v N_v + a_{max} N_h}\ bites \cdot day^{-1} \cdot human^{-1}$$

Similarly, $E_v$ represents the average number of effective encounters generated by each mosquito per unit time.

$$E_v = \frac{E_T}{N_v} = \frac{\sigma_v \; a_{max} \, N_h}{\sigma_v N_v \; + \; a_{max} N_h} \; bites \cdot day^{-1} \; \cdot mosquito^{-1}$$

From these quantities, the force of infection on humans is defined as

$$\lambda_h = E_h \cdot \beta_{hv} \cdot \frac{I_v}{N_v}$$

where $\beta_{hv}$ is the probability of transmission from mosquito to a human given contact, and $\frac{I_v}{N_v}$ is the proportion of infectious mosquitoes. Similarly, the force of infection on mosquitoes is

$$\lambda_v = E_v \cdot \beta_{vh} \cdot \frac{I_h}{N_h}$$

where $\beta_{vh}$ is the probability of transmission from human to a mosquito given contact and $\frac{I_h}{N_h}$ is the proportion of infectious humans. Under this formulation, transmission intensity emerges from the interaction among mosquito biting demand, human biting availability, and infection prevalence in the opposite population, rather than from a fixed categorical transmission regime.

#### S3. Event-driven implementation and state transitions

Model dynamics were represented through a combination of random events and fixed timers. Human infection, mosquito infection, human recovery, and mosquito death were treated as random events. In contrast, parasite development within hosts and vectors was modeled using individual timers set at the moment of inoculation or acquisition.

In humans, newly inoculated parasites require a fixed maturation time ($\alpha_h$), before producing gametocytes and becoming infectious. In mosquitoes, parasites similarly require a fixed maturation time ($\alpha_v$), before developing into transmissible sporozoites. These maturation steps were modeled deterministically, allowing the model to separate parasite acquisition from the onset of infectiousness. Because the main objective of this study is to analyze the evolution of parasite population genetic structure, the model was framed under an endemic transmission setting in which infection establishment in humans was assumed to be somewhat less efficient, while infectious *Anopheles* vectors remained effective at transmitting the parasite.

Each inoculation event was tracked independently, allowing multiple infections to coexist within the same host or vector. Thus, an individual could carry mature parasites from a previous infection while simultaneously harboring newly acquired immature parasites from a subsequent inoculation. State changes were resolved through an event queue, in which the next scheduled event corresponded to the earliest pending transition across all individuals. This event-driven structure preserved temporal resolution while accommodating repeated exposure, overlapping infections, and developmental delays.

| Symbol | Units | Value / Range | Description | Source |
|---|---|---|---|---|
| $N_h$ | humans | 50 | Human population size | - |
| $m$ | mosquitoes/human | 5 | Mosquito-to-human ratio | - |
| $N_v$ | mosquitoes | 250 | Total mosquito population | - |
| $\sigma_v$ | bites·mosquito$^{-1}$·day$^{-1}$ | $(3.1)^{-1}$ | Mosquito biting demand, related to the gonotrophic cycle | (Rúa, 2005) |
| $a_{max}$ | bites·human$^{-1}$·day$^{-1}$ | 0.1–3.0 | The maximum bites a human can receive per unit time | (Chitnis, 2008) |
| $\beta_{hv}$ | — | 0.2 | Transmission probability from mosquito to human | (Aleshnick, 2020) |
| $\beta_{vh}$ | — | 0.07 | Transmission probability from human to mosquito | (Churcher, 2015) |
| $\gamma_h$ | days | 289 | Human recovery time | (Chitnis, 2008) |
| $\delta$ | days | 30 | Mosquito lifespan | (Osoro, 2022) |
| $\alpha_h$ | days | 11 | Gametocyte maturation time in humans | (Alemayehu, 2023) |
| $\alpha_v$ | days | 14 | Sporozoite maturation time in mosquitoes | (Alemayehu, 2023) |
| $\pi_0$ | % | 10, 30, 50, 75, 90 | Initial genetic diversity levels | - |
| $L$ | nucleotide sites | 75 | Haplotype length | - |

**Table S1. Parameter definitions, explored ranges, and sources used in the transmission-genetics model.** Baseline values were used for primary simulations, whereas explored ranges were included to assess biologically plausible parameter variation.

#### S4. Parasite representation and initialization of genetic diversity

Parasite populations were represented by haplotypes defined as fixed-length positional marker sequences. In the simulations presented here, each haplotype consisted of $L = 75$ positions, a length that proved sufficient to prevent saturation in the number of simulated haplotypes. The model allows repeated infections by the same haplotype, as well as simultaneous infection by multiple distinct haplotypes within the same host or vector.

Initial parasite diversity was established by defining a starting haplotype pool that matched a predetermined level of genetic diversity, denoted as $\pi_0$, and five levels of initial diversity were considered (Table S1). For each scenario, haplotypes from the starting pool were assigned to infected individuals at the start. When the initial haplotype pool contained relatively few haplotypes, these were evenly distributed across the infected population to prevent strong compositional bias at the beginning of the simulation.

Baseline simulations were initialized with a mosquito-to-human ratio of 5:1, with all humans and mosquitoes initially infected and each infection monoclonal. This initialization strategy allowed the simulations to isolate the effects of predefined starting diversity on the subsequent emergence of mixed infections, recombination, and population-level genetic structure over time.

#### S5. Recombination model

Parasite recombination was modeled during the mosquito stage. Recombination could occur when a mosquito acquired multiple haplotypes in a single infectious blood meal, allowing different parasite genotypes to enter the sexual stage within the vector. Mixed infections in mosquitoes, therefore, provide the opportunity for recombinant offspring to be produced. The number of oocysts formed in such events was modeled as a Poisson random variable with a mean of 2 (McKenzie, 2002; Stopard, 2021). For each mixed infection, recombinant offspring were generated by recombining parental haplotypes through crossover events across the genome. In the reported simulations, recombination breakpoints were sampled uniformly across the 75-position genome, implying no recombination hotspots. Although the model framework can include non-uniform recombination landscapes, such as hotspot-based maps, these were not used in the current analyses. This recombination procedure allowed the model to distinguish between recombination events that merely reshuffled existing variation and those that generated haplotypes not previously present in the circulating parasite population.

#### S6. Outcome metrics

Model behavior was characterized using a set of epidemiological and genetic summary metrics. Epidemiological outcomes comprised infection prevalence, the prevalence of monoclonal and polyclonal infections in humans and mosquitoes, force of infection, and multiplicity of infection (MOI) for each host type. Genetic outcomes comprised population nucleotide diversity ($\pi$), normalized Shannon diversity, the number of haplotypes, and the effective recombination ratio.

| Category | Metric | Definition |
|---|---|---|
| Epidemiological | Infection prevalence | Proportion of sampled hosts that are infected, computed as the number of infected individuals divided by the total number of individuals sampled. |
| Epidemiological | Monoclonal infection prevalence | Proportion of infections containing exactly one haplotype, computed as the number of monoclonal infections divided by the total number of infections. |
| Epidemiological | Polyclonal infection prevalence | Proportion of infections containing more than one haplotype, computed as the number of polyclonal infections divided by the total number of infections. |
| Epidemiological | Multiplicity of infection (MOI) | Number of haplotypes carried by an infected individual, summarized separately across infected humans and infected mosquitoes. |
| Genetic | Nucleotide diversity ($\pi$) | Average pairwise genetic difference between haplotypes, weighted by haplotype frequencies, computed as $\pi = 2\sum_{i<j} \pi_i \pi_j d_{ij}$ where $\pi_i$ and $\pi_j$ are the relative frequencies of haplotypes $i$ and $j$. Then, $d_{ij} = \frac{1}{L} \sum_{l=1}^{L} 1 \{h_{il} \neq h_{jl}\}$ is the normalized Hamming distance between the two haplotypes across the $L$ nucleotide positions. |
| Genetic | Normalized Shannon diversity | Diversity of haplotype frequencies normalized by the number of observed haplotypes, computed as $H_{\text{norm}} = -\frac{\sum_i p_i \log(p_i)}{\log(S)}$ where $p_i$ is the frequency of haplotype $i$ and $S$ is the number of observed haplotypes. When $S \leq 1$, $H_{\text{norm}}$ was set to 0. |
| Genetic | Number of haplotypes | Total number of distinct haplotypes observed in the circulating parasite population during the time window considered. |
| Genetic | Effective recombination ratio | Fraction of infection events in a given time interval that generate at least one novel haplotype through recombination, computed as: $R_{\text{eff}}(t) = \frac{N_{\text{novel}}(t)}{N_{\text{infection}}(t)}$ where $N_{\text{novel}}(t)$ is the number of infection events in time interval $t$ that produce at least one novel haplotype through recombination, and $N_{\text{infection}}(t)$ is the total number of infection events in the same interval. |

**Table S2. Definitions of epidemiological and genetic summary metrics used to quantify model behavior.** The table summarizes the epidemiological and genetic outcome measures used in the analysis, together with their mathematical definitions where applicable.

### S7. Supplementary Figures

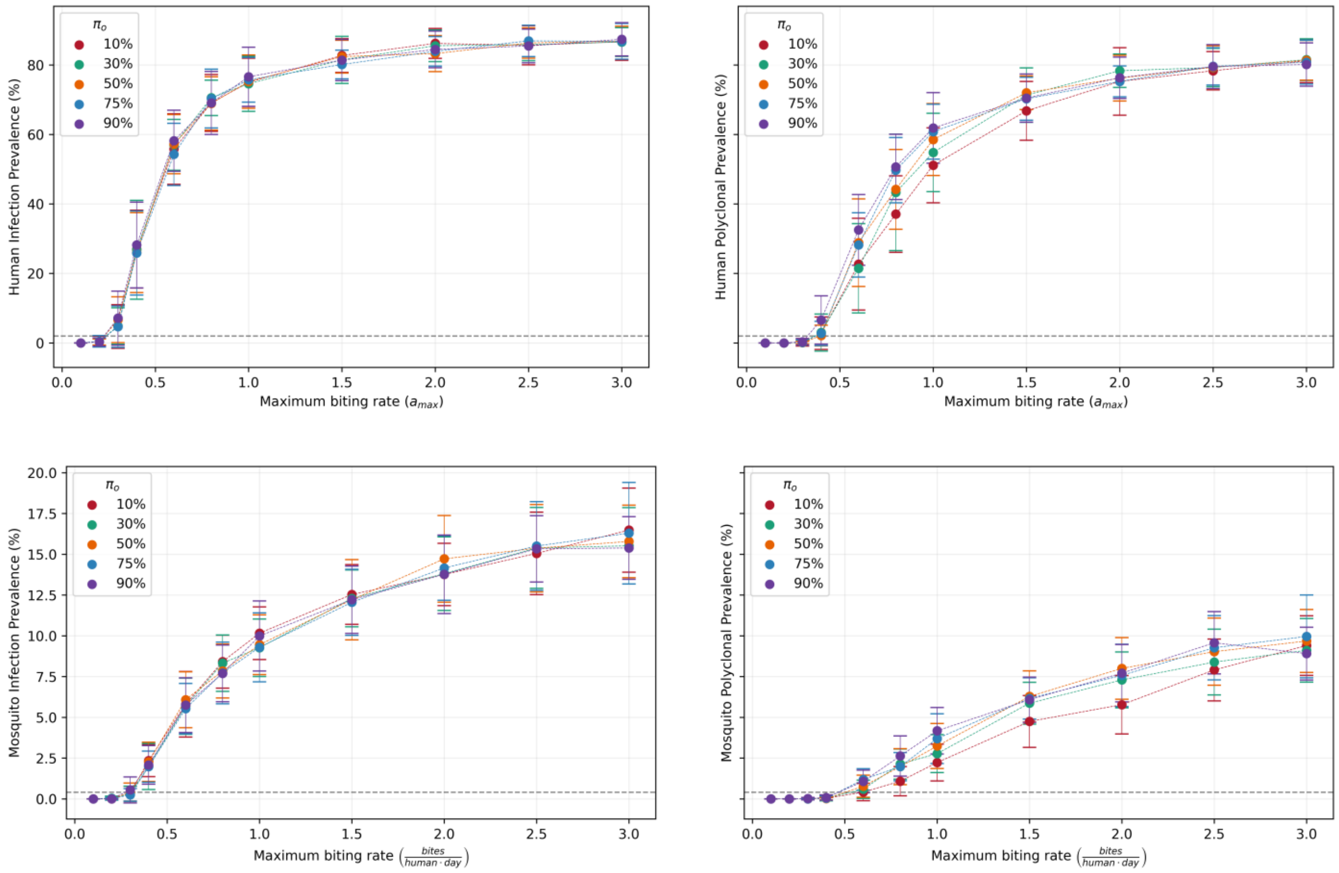


**Figure S2. Steady-state infection prevalence in humans and mosquitoes and prevalence of polyclonal infections as functions of the maximum biting rate.** Colors indicate different levels of initial parasite genetic diversity, and each point represents the mean across simulations for a given parameter combination. The dashed horizontal line marks the one-infected-human threshold; values above this line indicate persistence of infection in at least one human at steady state.

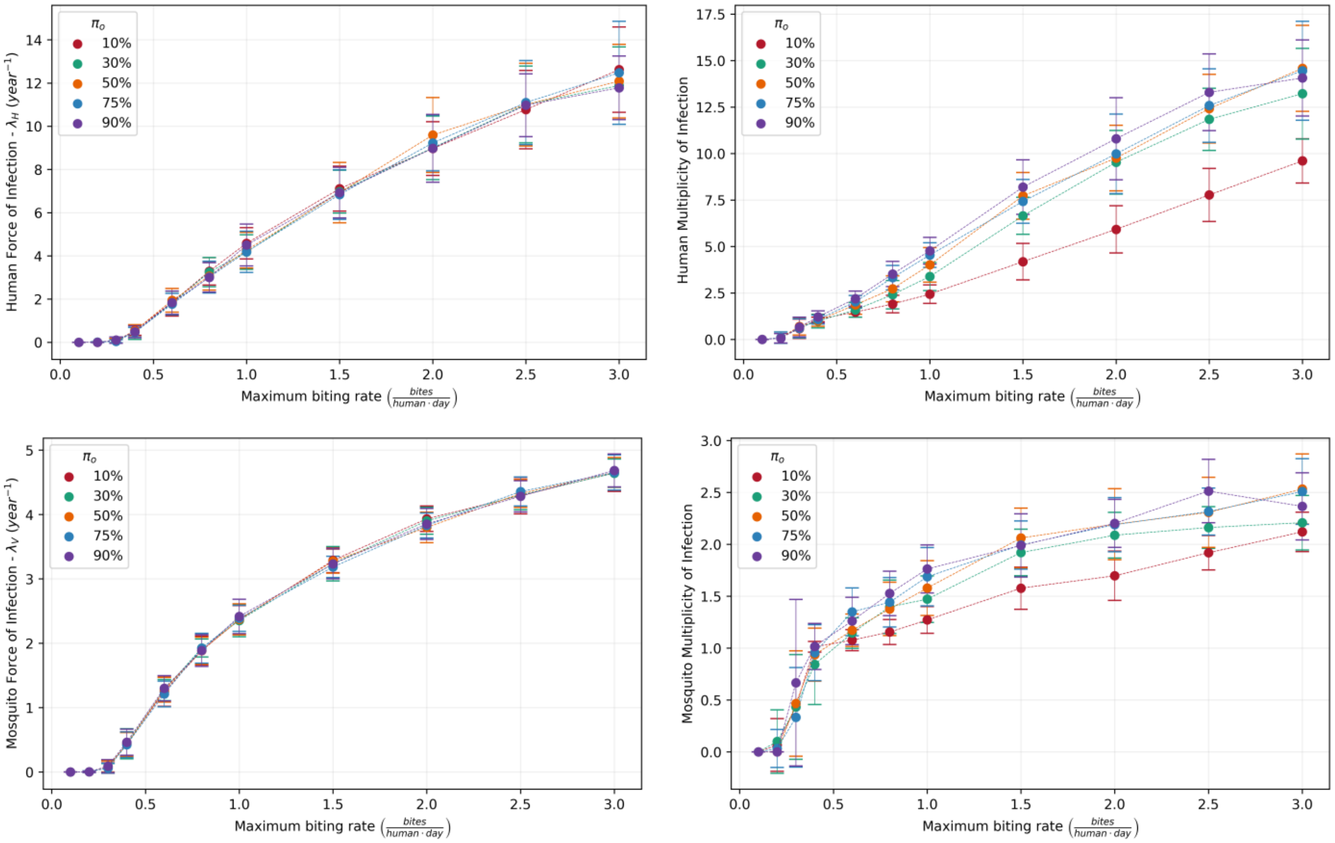


**Figure S3. Steady-state force of infection (FOI) and mean multiplicity of infection (MOI) as functions of the maximum biting rate in human and mosquito populations.** Colors indicate different levels of initial parasite genetic diversity, and each point represents the mean across simulations for each parameter combination.

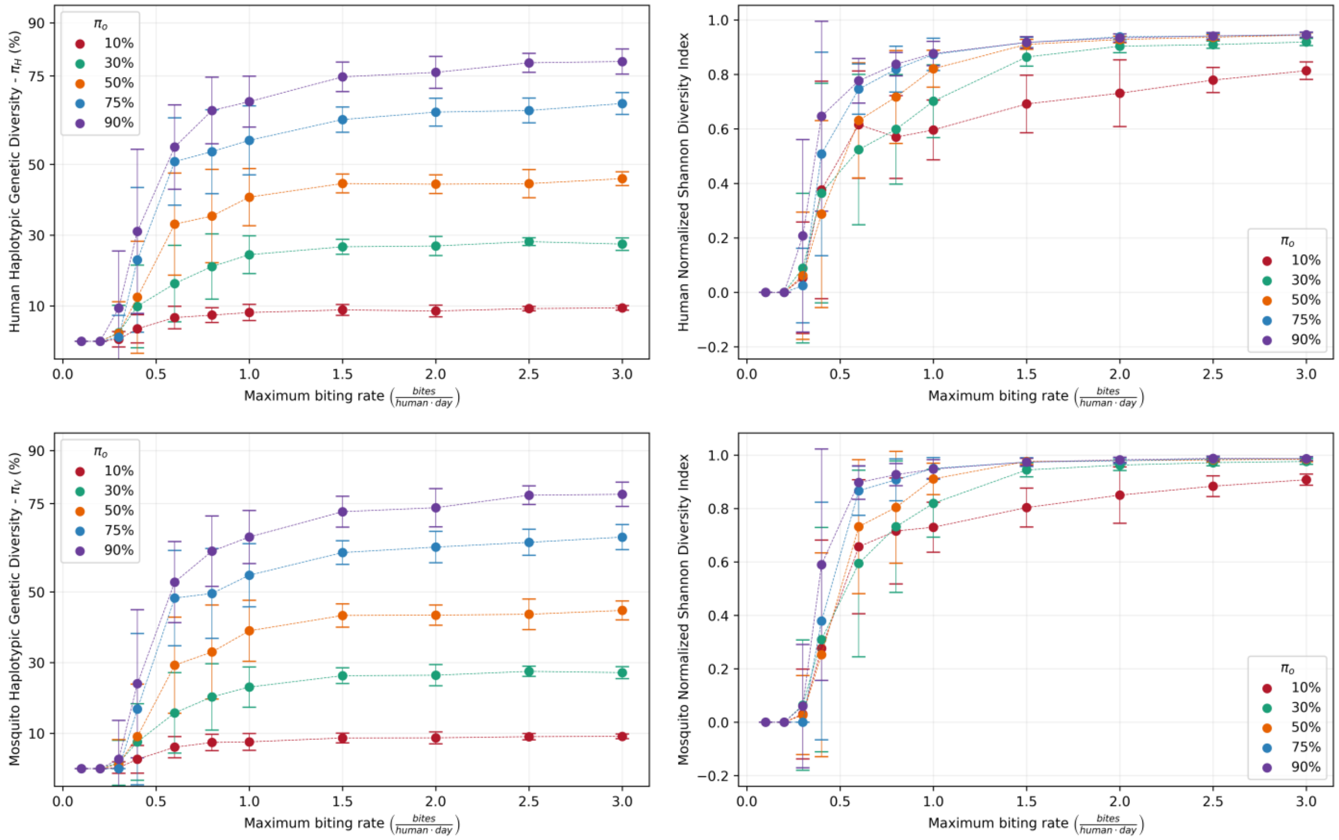


**Figure S4. Steady-state haplotypic genetic diversity and normalized Shannon index as functions of the maximum biting rate in human and mosquito populations.** Colors indicate different levels of initial parasite genetic diversity, and curves represent the mean across simulations for each parameter combination.

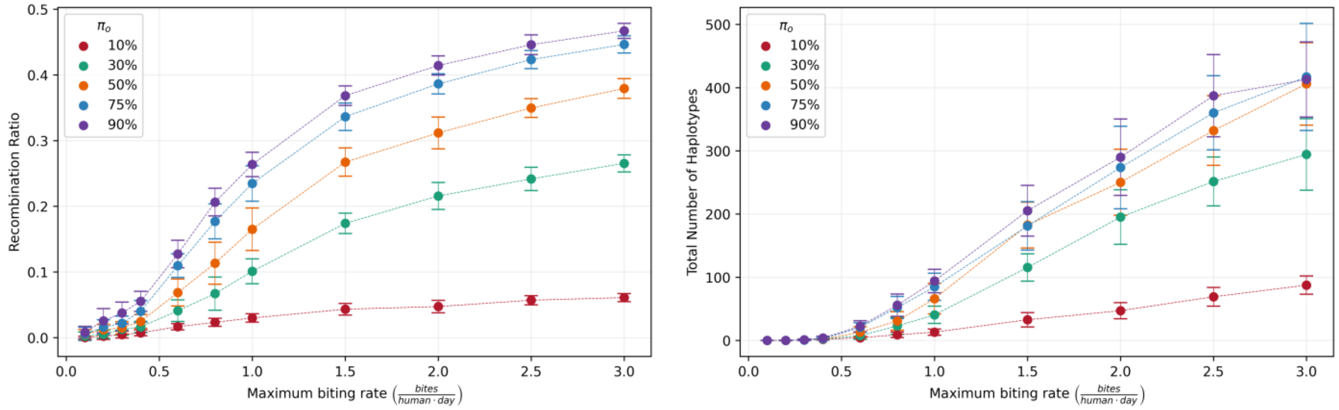


**Figure S5. Steady-state recombination ratio and total number of haplotypes as functions of the maximum biting rate in the parasite population**. Colors indicate different levels of initial parasite genetic diversity, and curves represent the mean across simulations for each parameter combination.

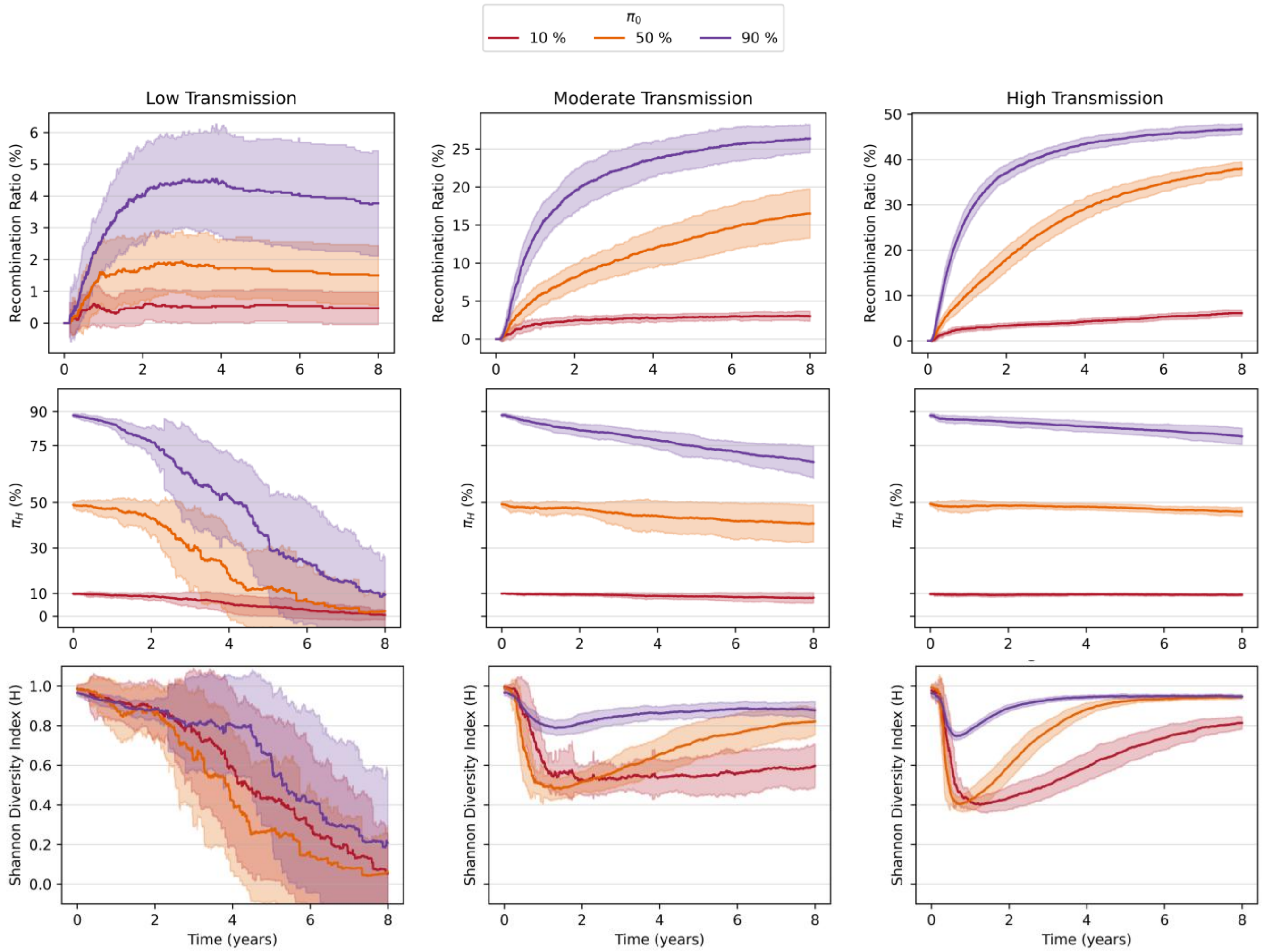


**Figure S6.** Temporal dynamics of the recombination ratio, human haplotypic genetic diversity ($\pi_H$) and normalized Shannon diversity index under low ($a_{\max} = 0.3$), moderate ($a_{\max} = 1.0$), and high ($a_{\max} = 3.0$) transmission scenarios. Columns indicate transmission intensity, whereas colors denote different initial levels of parasite genetic diversity. Curves show the mean across simulations, and shaded regions indicate variability across simulations.

#### S8. Demographic sensitivity to human population size and mosquito-to-human ratio

To assess whether the steady-state patterns described in the main analysis depended strongly on the baseline demographic setting ($N_h = 50$ humans and $m = 5$ mosquitoes per human), we repeated simulations across three human population sizes ($N_h = 30, 50, 70$) and three mosquito-to-human ratios ($m = 3, 5, 7$), while retaining representative low, intermediate, and high values of both initial genetic diversity ($10\%, 50\%, 90\%$) and maximum biting rate ($0.3, 1.0, 3.0$). The resulting heatmaps therefore separate two distinct demographic manipulations: variation in contact structure through $m$, and variation in total system size through $N_h$.

This distinction is important for interpretation. Under the harmonic-mean encounter formulation, varying $m$ modifies how encounters are distributed between hosts and vectors, whereas varying $N_h$ at fixed $m$ mainly changes the total number of available encounters, and with it the amount of stochastic loss, persistence, and recombinant retention in the system. The figures below therefore should not be read as a search for identical heatmaps across all demographic combinations. Instead, they test whether the main qualitative relationships between transmission, mixed infection, recombination, and diversity are retained once both contact structure and system size are allowed to vary.

Under the biting formulation used in Supplementary Section S2, the total number of effective encounters is given by:

$$E_T = \frac{(a_{max}\, N_h\, \sigma_v\, N_v)}{(a_{max}\, N_h + \sigma_v\, N_v)}$$

Thus, the per-human and per-mosquito encounter rates where $m = N_v/N_h$ are given by:

$$E_h = \frac{(a_{max}\, \sigma_v\, m)}{(a_{max} + \sigma_v\, m)} \qquad E_v = \frac{(a_{max}\, \sigma_v)}{(a_{max} + \sigma_v m)}$$

This decomposition makes clear that varying the mosquito-to-human ratio changes the contact structure itself, whereas varying the number of humans at fixed $m$ primarily rescales system size and the total number of encounters. As a result, identical heatmaps are not expected when $m$ changes. Epidemiological metrics that depend strongly on mosquito infection should be especially sensitive to $m$, whereas richness and diversity metrics may also respond to $N_h$ through changes in drift, extinction, and the persistence of recombinant haplotypes. The explored gradient therefore samples both contact-structure effects and system-size effects, but it does not fully compensate for increases in humans with decreases in mosquitoes; true compensation would require holding $E_h$ or $E_v$ approximately constant by jointly adjusting $m$ and $a_{max}$.

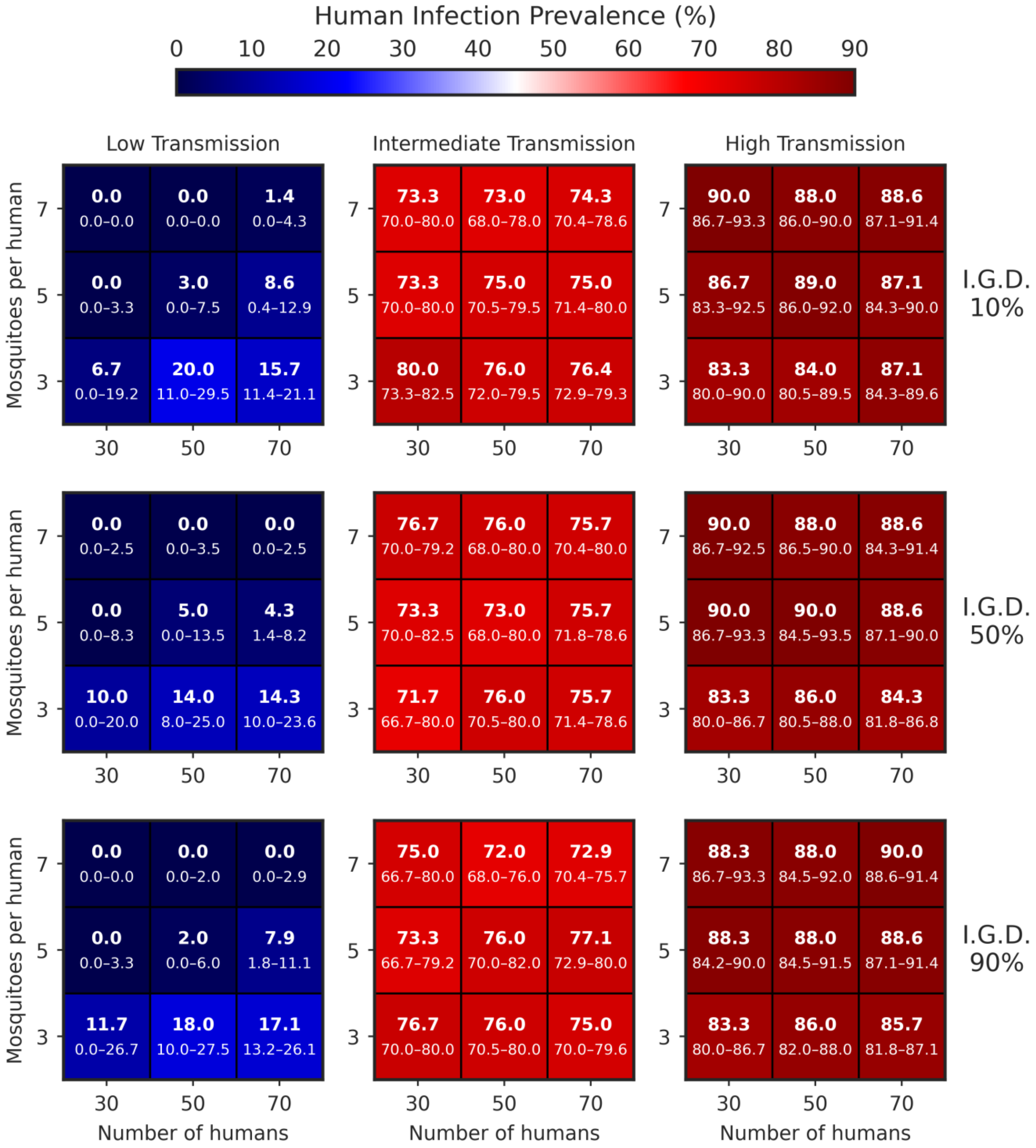


**Figure S7. Human infection prevalence at steady state across demographic sensitivity simulations.** In each cell, the larger number indicates the median across replicate simulations, whereas the smaller numbers indicate the interquartile range (25th–75th percentiles).

Each heatmap cell reports the median steady-state human infection prevalence across replicate simulations, with the interquartile range shown below the median. Across all initial diversity levels, prevalence increases strongly from low to intermediate and high transmission. Within each transmission regime, variation across the mosquito-to-human ratio is generally stronger than variation across human population size, indicating that contact structure has a larger effect on epidemiological burden than system size alone. The fact that prevalence does not remain constant across heatmaps is expected under the model formulation, because changing $m$ alters vector-host encounters rather than simply rescaling population size.

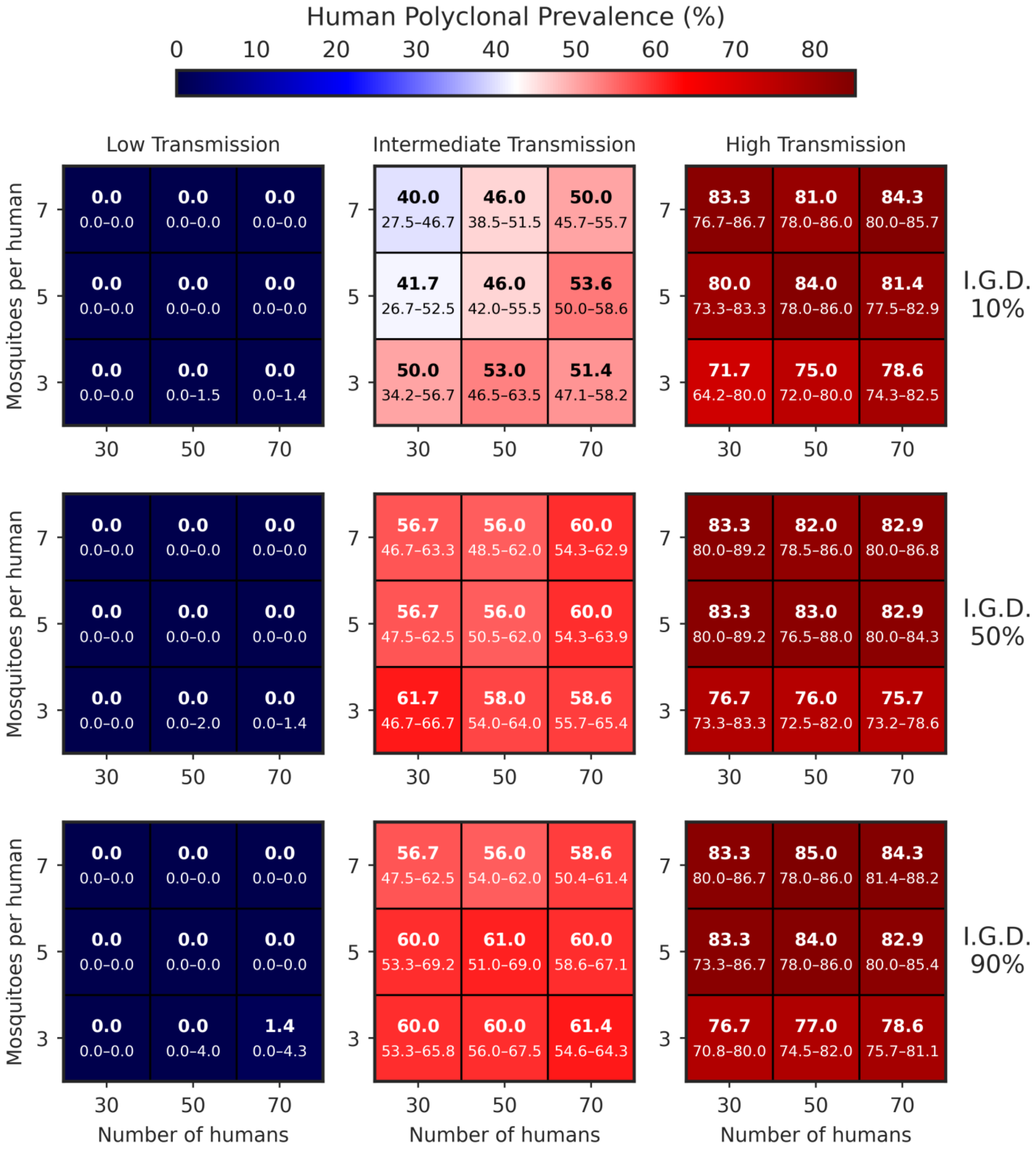


**Figure S8. Human polyclonal infection prevalence at steady state across demographic sensitivity simulations.** In each cell, the larger number indicates the median across replicate simulations, whereas the smaller numbers indicate the interquartile range (25th–75th percentiles).

Figure S8 shows the fraction of infected humans carrying more than one haplotype. Human polyclonality is near zero in low-transmission settings and rises sharply under intermediate and high transmission, consistent with the idea that mixed infection is an intermediate step linking transmission to recombination. As in Figure S7, differences across the mosquito-to-human ratio are more pronounced than differences across human population size, reinforcing that within-host complexity is governed primarily by encounter structure rather than by total population size alone.

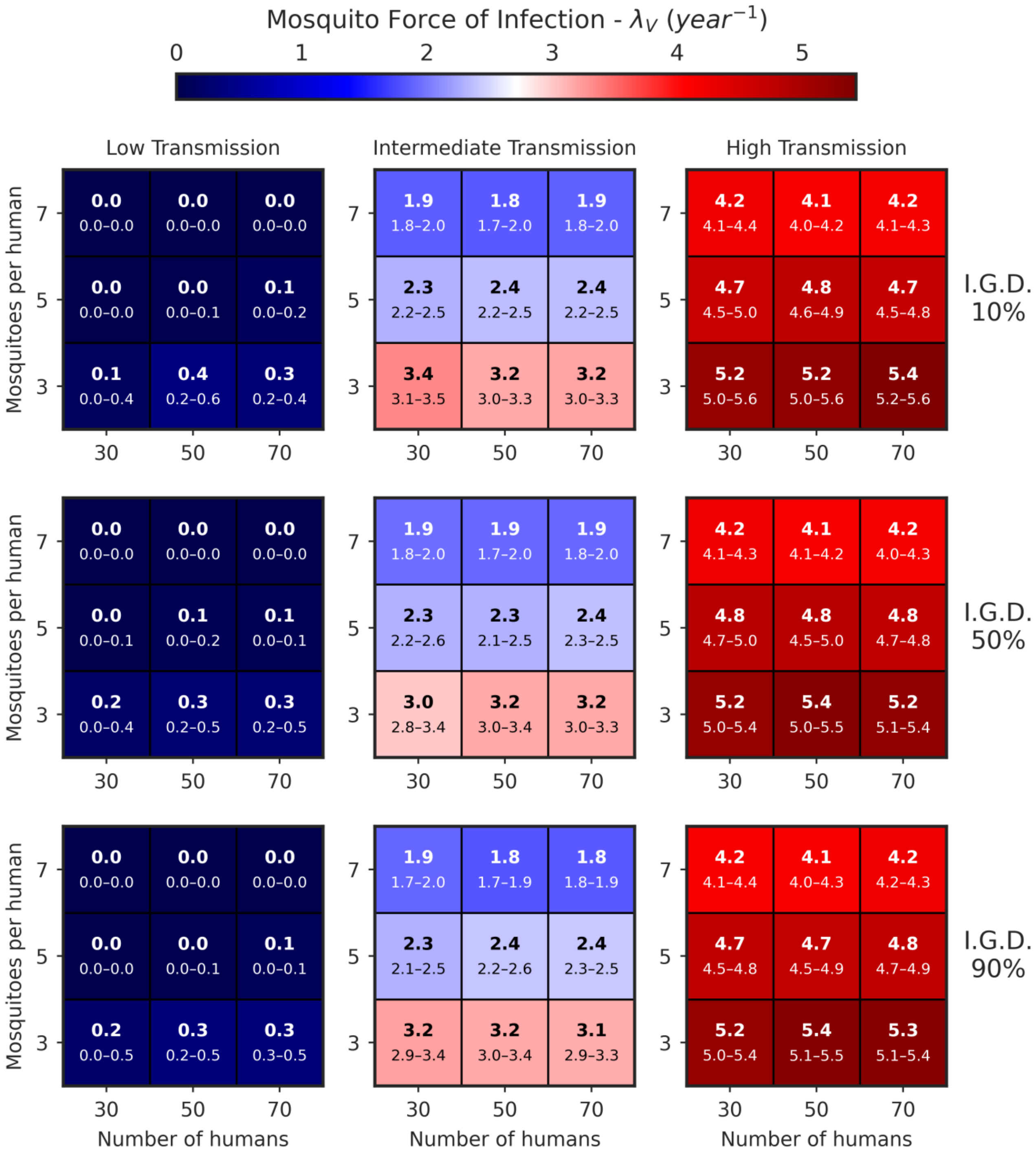


**Figure S9. Mosquito force of infection at steady state across demographic sensitivity simulations.** In each cell, the larger number indicates the median across replicate simulations, whereas the smaller numbers indicate the interquartile range (25th–75th percentiles).

This figure provides the most direct confirmation of the contact formulation used in the model. Because the per-mosquito encounter rate decreases as the mosquito-to-human ratio increases, the mosquito force of infection is expected to be most sensitive to $m$ and only weakly sensitive to $N_h$ when $m$ is held fixed. The heatmaps are consistent with that expectation: within a given transmission regime, $\lambda_v$ changes systematically across the mosquito-to-human gradient, while the effect of human population size is comparatively modest.

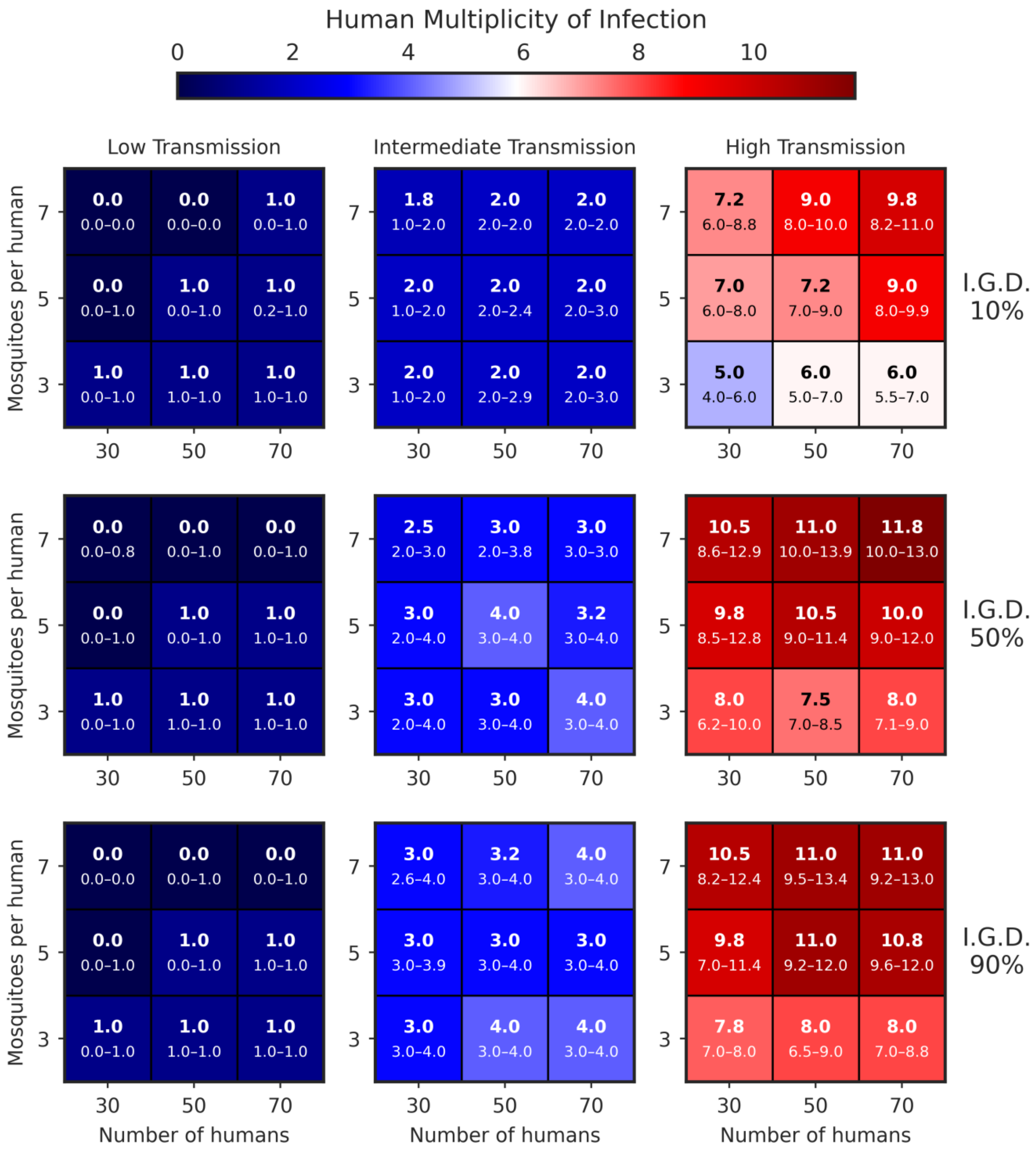


**Figure S10. Human multiplicity of infection at steady state across demographic sensitivity simulations.** In each cell, the larger number indicates the median across replicate simulations, whereas the smaller numbers indicate the interquartile range (25th–75th percentiles).

Human multiplicity of infection (MOI) rises across the transmission gradient and also increases with initial diversity, reflecting greater opportunities for distinct haplotypes to co-occur within the same host. The demographic sensitivity runs show that MOI responds more strongly to the mosquito-to-human ratio than to human population size, especially once transmission is high enough for mixed infection to become common. This is important because MOI is the proximal within-host mechanism that enables recombination and therefore mediates the downstream genetic effects seen in later figures.

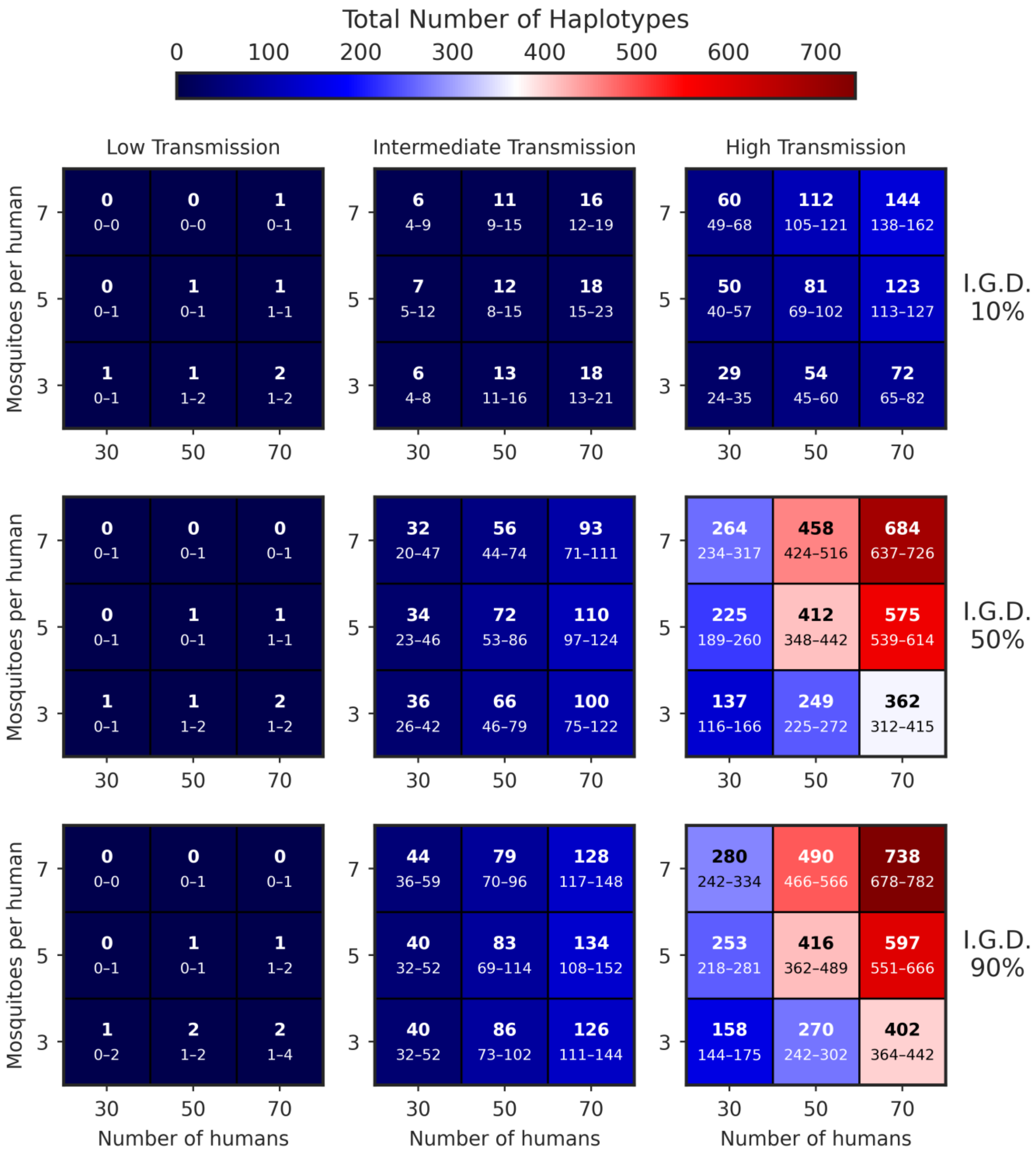


**Figure S11. Total number of circulating haplotypes at steady state across demographic sensitivity simulations.** In each cell, the larger number indicates the median across replicate simulations, whereas the smaller numbers indicate the interquartile range (25th–75th percentiles).

Unlike prevalence and force-of-infection metrics, haplotype richness shows a clearer dependence on both contact structure and system size. At intermediate and high transmission, increasing human population size can substantially increase the number of circulating haplotypes, especially when initial diversity is already moderate or high. This is expected as larger system produces more total encounters and therefore more opportunities for rare or recombinant haplotypes to be generated and to persist, even when per-capita transmission conditions are similar.

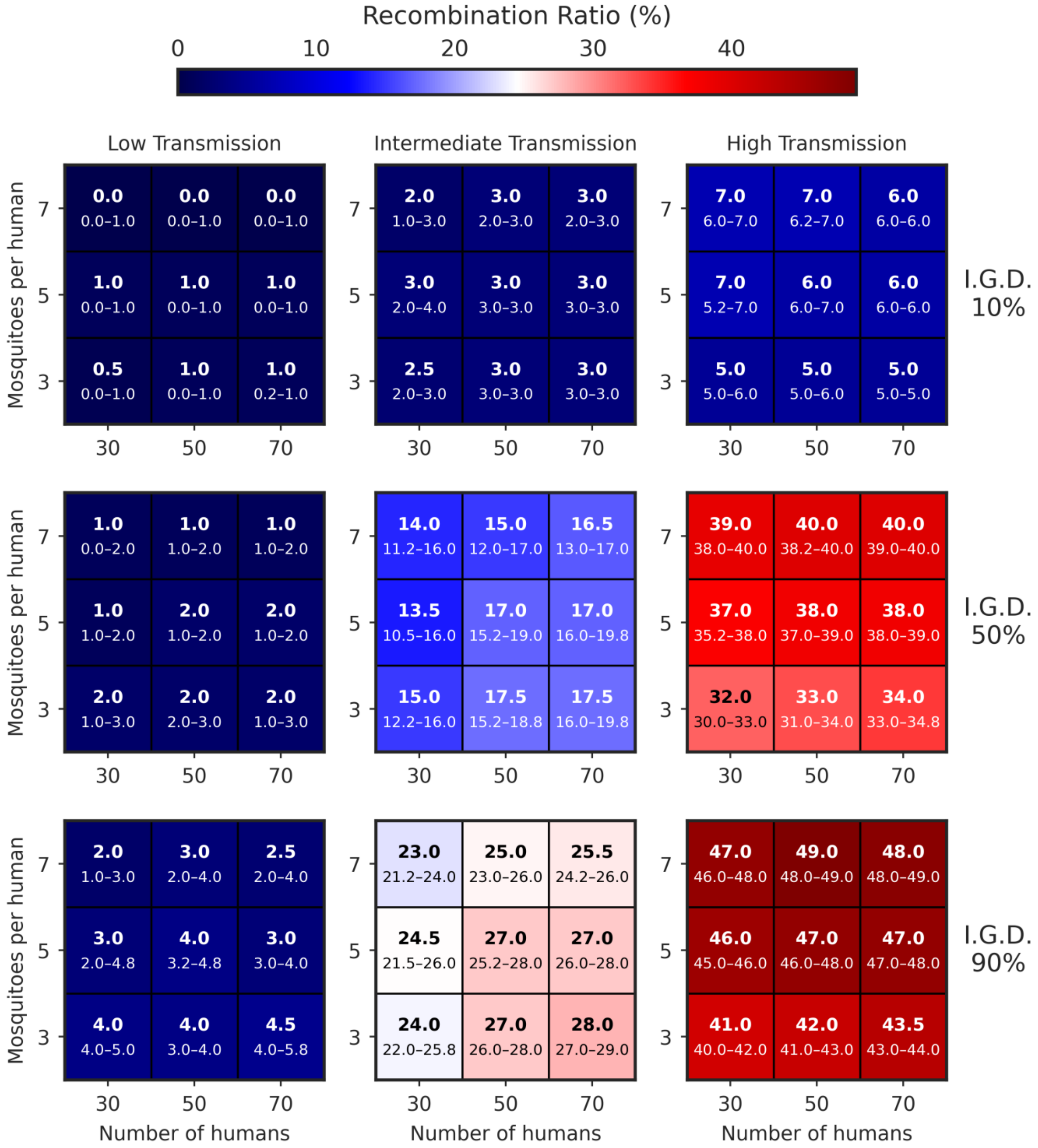


**Figure S12. Effective recombination ratio at steady state across demographic sensitivity simulations.** In each cell, the larger number indicates the median across replicate simulations, whereas the smaller numbers indicate the interquartile range (25th–75th percentiles).

The effective recombination ratio remains low under low transmission and increases under intermediate and high transmission, with especially strong increases at higher initial diversity. The demographic gradients show that recombination is shaped jointly by contact structure and standing diversity: changing the mosquito-to-human ratio alters how efficiently mixed infections are sustained, while larger systems can retain more recombinant products once they are generated. Thus, recombination-related outcomes are not purely a function of biting intensity, but also of the demographic context in which coinfection occurs.

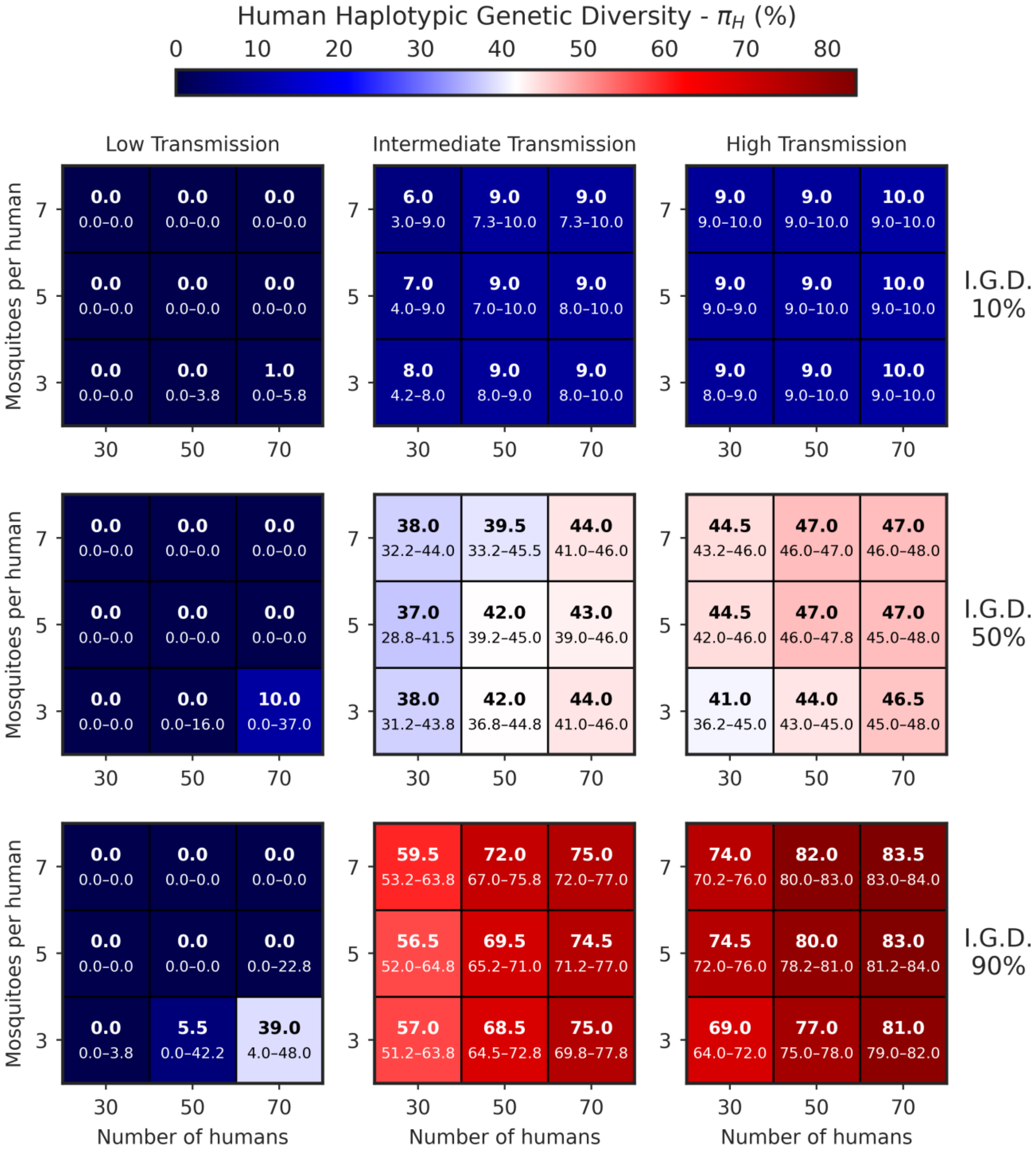


**Figure S13. Human haplotypic genetic diversity at steady state across demographic sensitivity simulations.** In each cell, the larger number indicates the median across replicate simulations, whereas the smaller numbers indicate the interquartile range (25th–75th percentiles).

Human haplotypic genetic diversity is especially sensitive to the interaction between initial diversity, transmission intensity, and demographic scale. A particularly visible pattern occurs when the mosquito-to-human ratio is low: increasing the number of humans can substantially raise the final diversity level, especially at intermediate and high initial diversity. This reflects a system-size effect rather than a change in per-capita contact alone, because larger systems are less prone to stochastic loss of lineages and better able to retain haplotypic variation once transmission and recombination are active.

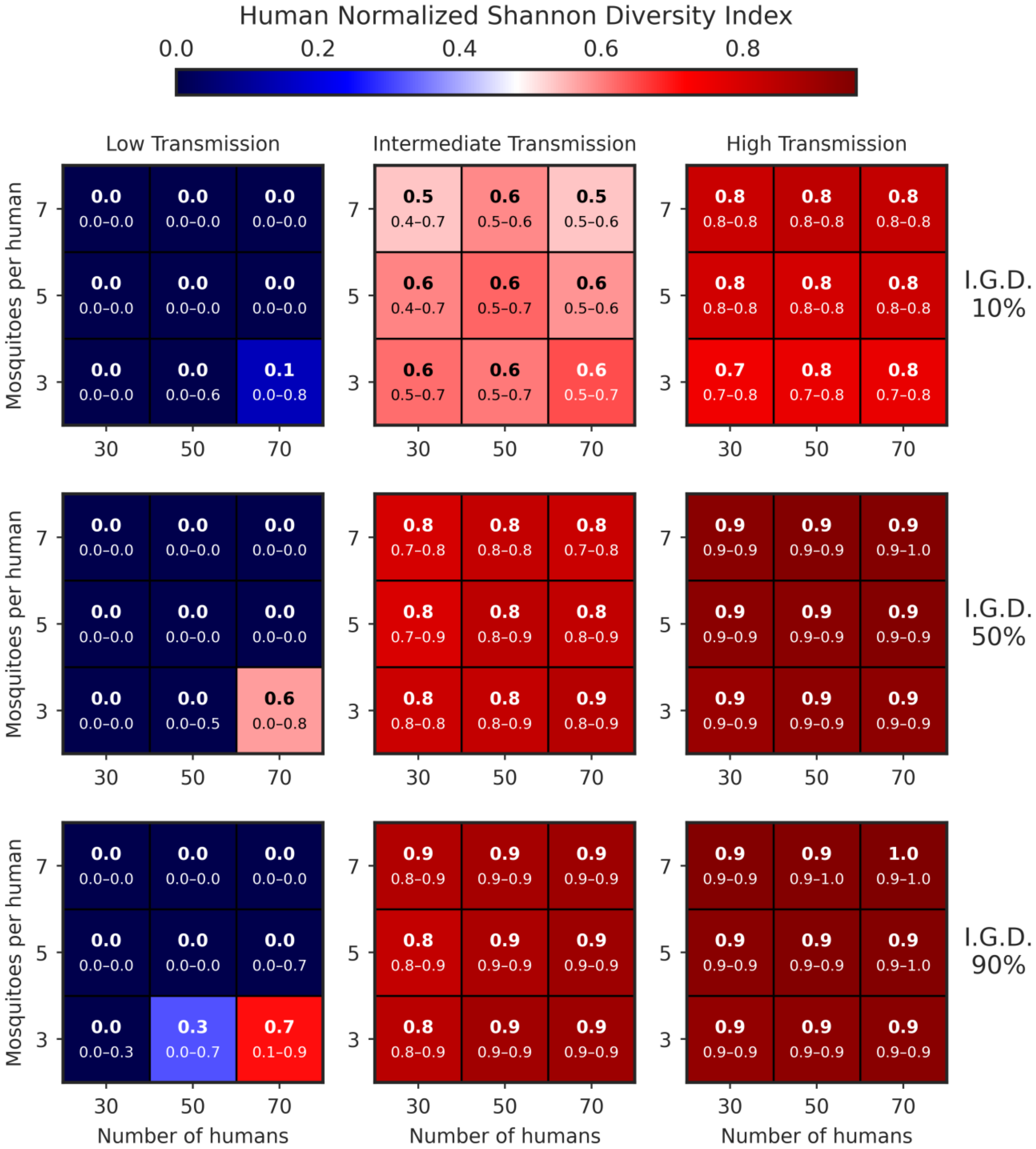


**Figure S14. Human normalized Shannon diversity index at steady state across demographic sensitivity simulations.** In each cell, the larger number indicates the median across replicate simulations, whereas the smaller numbers indicate the interquartile range (25th–75th percentiles).

The normalized Shannon index complements Figure S13 by showing how evenness responds to the same demographic gradients. At low transmission, evenness collapses toward zero across most demographic settings, indicating domination by one or a few haplotypes. At intermediate and high transmission, evenness increases sharply, but its final value remains influenced by both the mosquito-to-human ratio and total system size when diversity is still being assembled. Together, we show that genetic outcomes are shaped not only by whether transmission is high or low, but also by the demographic conditions that determine how efficiently diversity can be maintained once generated.

Combined, these sensitivity analyses show that the baseline conclusions of the model are robust, but not invariant, to demographic scaling. Varying the mosquito-to-human ratio changes the encounter structure itself and therefore alters mosquito acquisition, mixed infection, and downstream recombination. Varying the number of humans at fixed $m$ changes the size of the transmission system and therefore the amount of stochastic loss or retention of haplotypes. For that reason, epidemiological metrics are expected to be most sensitive to $m$, whereas genetic richness and evenness can retain additional dependence on $N_h$. This distinction is precisely what makes the sensitivity analysis informative: it shows that the main transmission-genetic relationships remain qualitatively intact, while also clarifying which outcomes are most responsive to contact structure and which remain influenced by system size.
