## Supplementary material for "From low to high transmission: Diversity-dependent responses of *Plasmodium falciparum* population structure to transmission intensity": License Figure S1

49 Spadina Ave. Suite 200
Toronto ON M5V 2J1 Canada
www.biorender.com

### Confirmation of Publication and Licensing Rights - Open Access

**April 1st, 2026**

| | |
|---|---|
| ***Subscription Type:*** | *Institution - Academic* |
| ***Agreement number:*** | *HJ29JLWOYE* |
| ***Publisher Name:*** | *The Royal Society* |
| ***Figure Title:*** | *Extended state-transition structure for human and mosquito infections.* |
| ***Citation to Use:*** | *Created in BioRender. Suarez Salazar, C. (2026) https://BioRender.com/xxj2op8* |

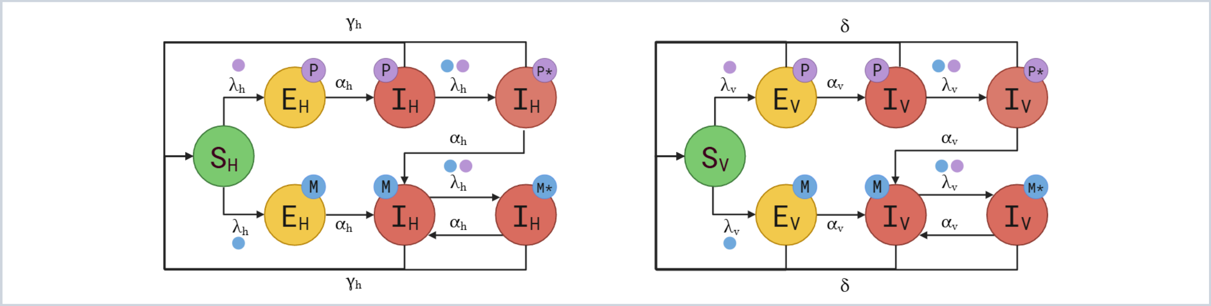


*For any questions regarding this document, or other questions about publishing with BioRender, please refer to our BioRender Publication Guide, or contact BioRender Support at.*
